# The fluctuations of alpha power: bimodalities, connectivity, and neural mass models

**DOI:** 10.1101/2025.03.04.641398

**Authors:** Jesús Cabrera-Álvarez, Alberto del Cerro-Léon, Blanca P. Carvajal, Martín Carrasco-Gómez, Christoffer G. Alexandersen, Ricardo Bruña, Fernando Maestú, Gianluca Susi

## Abstract

The alpha rhythm is a hallmark of electrophysiological resting-state brain activity, that serves as a biomarker in health and disease. Alpha power is far from uniform over time, exhibiting dynamic fluctuations. The likelihood of those power values can be captured by a decreasing exponential function, that in certain cases becomes bimodal. While alpha rhythm is usually evaluated through the averaged power spectra across entire recordings, its dynamic fluctuations have received less attention. In this study, we investigate the dynamic nature of alpha power, its relationship with functional connectivity (FC) within the default mode network (DMN), and the ability of the Jansen-Rit (JR) neural mass model to replicate these fluctuations. Using MRI and MEG data from 42 participants in resting state with eyes-closed and eyes-open, we evaluated the shape of the exponential distributions for alpha power fluctuations, and their relationship with other spectral variables as frequency, power and the aperiodic exponent. Additionally, we assessed the temporal relationship between alpha power and FC using phase-based (ciPLV) and amplitude-based (cAEC) metrics. Finally, we employed diffusion-weighted MRI to construct brain network models incorporating JR neural masses to reproduce and characterize alpha fluctuations. Our results indicate that alpha power predominantly follows unimodal exponential distributions, with bimodalities associated to high-power in posterior regions. FC analyses revealed that ciPLV and cAEC were directly correlated with alpha power within the DMN in alpha and beta bands, whereas only theta-band ciPLV showed an inverse relationship with alpha power. JR model simulations suggested that post- supercritical fixed points better replicated alpha power fluctuations compared to limit cycle parameterizations and pre- saddle node fixed points. These results deepen our understanding of the dynamics of alpha rhythm and its intricate relationship with FC patterns, offering novel insights to refine biologically plausible brain simulations and advance computational models of neural dynamics.

## Introduction

Alpha oscillations are a fundamental brain rhythm characterized by electrophysiological fluctuations at around 10 cycles per second. First identified by Hans Berger (Berger, 1929), these oscillations have emerged as a crucial biomarker both in health and disease (Ippolito et al., 2022; Babiloni et al., 2025; Zhang et al., 2024; Ye et al., 2022). Alpha activity manifests most prominently during periods of resting state with eyes closed (Barry et al., 2007; Wan et al., 2018; Ingram et al., 2024), but its power also appears modulated in task-related contexts such as working memory (Jensen, 2002; Tuladhar et al., 2007; Bonnefond and Jensen, 2012; Wianda and Ross, 2019; Palva and Palva, 2007) and attentional tasks (Sauseng et al., 2005; Kelly et al., 2006; Foxe and Snyder, 2011).

Alpha has been conceptualized both as an active state of cortical inhibition that avoids irrelevant stimuli during highly demanding cognitive tasks (Jensen, 2024) and as a passive state of cortical idling (Pfurtscheller et al., 1996). These two theories have been supported by multiple lines of evidence including the observation of anti-correlations between BOLD signals and alpha power (Goldman et al., 2002; Ritter et al., 2008; Liu et al., 2014), the reductions in alpha amplitude within modality specific regions during cognitive processing -while increasing in task-irrelevant ones- (Pfurtscheller, 1992; Worden et al., 2000), and the enhancements of alpha during states of focused relaxation, such as mindfulness and meditation (Katyal and Goldin, 2021; Lomas et al., 2015).

Alpha oscillations fluctuate in amplitude over time, exhibiting waxing and waning dynamics. The likelihood of power values can be captured by a decreasing exponential function (Freyer et al., 2009), indicating a higher prevalence of low-power states with an exponentially decreasing likelihood for higher power values. Intriguingly, in some cases, these distributions appear bimodal, suggesting two distinct modes of alpha activity—one with higher and another with lower amplitude (Freyer et al., 2012). Such bimodalities have been interpreted within the framework of neuronal criticality reflecting transitions between two states (Freyer et al., 2012).

At the network level, alpha has been associated with the activation of the default mode network (DMN) from a time-averaged perspective. Electrophysiological studies have reported increased functional connectivity (FC) in the default mode network (DMN) associated to higher alpha power (Kim et al., 2023; Clancy et al., 2021). Other authors propose a more nuanced relationship, suggesting a direct association only in the eyes-open resting state (Mo et al., 2013), and highlighting region-dependent variations -some areas exhibiting a positive correlation with alpha power, while others show an inverse relationship (Bowman et al., 2017). However, whether there is a relationship between the fluctuations of alpha power and network dynamics remains an open question. Given the anti-correlations observed in previous studies between alpha power and BOLD signals, we hypothesize to find also an indirect relationship between alpha power and the activation of the DMN, taking into account the spatial complexity and the temporal dynamics of both variables.

Finally, neural mass models, such as the Jansen-Rit (JR) (Jansen and Rit, 1995), are able to reproduce the waxing-waning fluctuations of alpha, in this case, through the interaction of biologically plausible neuronal inhibitory and excitatory subpopulations. However, these dynamics can be obtained with different types of parameterizations, including limit cycles regimes in which the model autonomously oscillates, and fixed points regimes in which the model behaves as a damped oscillator (Grimbert and Faugeras, 2006; Spiegler et al., 2010). The accuracy of each of those regimes to resemble the empirical electrophysiological fluctuations of power observed in M/EEG remains to be studied.

In this study, we aim to advance our understanding of alpha oscillations through a comprehensive characterization of power fluctuations across regions and considering two brain states (resting with eyes-open and eyes-closed). Using MEG data, we will analyse the likelihood distribution of power values through time, identifying regions exhibiting bimodal distributions, and quantifying their prevalence. Additionally, we will examine the relationship between alpha power fluctuations and FC in the DMN. Finally, we will evaluate the JR model to reproduce the shape of the empirically observed alpha fluctuations, discussing the theoretical implications of different model parameterizations, and commenting on the mechanisms underlying alpha fluctuations.

## Materials and Methods

### Dataset

MRI (T1 and DWI) scans and MEG recordings were acquired from 42 healthy participants (20 females) with mean age 70.76 (std 4.65) from a dataset owned by the Centre for Cognitive and Computational Neuroscience, UCM, Madrid. MRI-T1 protocols were performed using a General Electric 1.5 tesla magnetic resonance scanner, using a high-resolution antenna and a homogenization PURE filter (fast spoiled gradient echo sequence, with parameters: repetition time/echo time/inversion time = 11.2/4.2/450 ms; flip angle = 12°; slice thickness = 1 mm, 256×256 matrix, and field of view = 256 mm).

DWI were acquired with a single-shot echo-planar imaging sequence with parameters: echo time/repetition time = 96.1/12,000 ms; NEX 3 for increasing the signal-to-noise ratio; slice thickness = 2.4 mm, 128×128 matrix, and field of view = 30.7 cm yielding an isotropic voxel of 2.4 mm. A total of 25 diffusion sampling directions were acquired with b-value = 900 s/mm2, plus one additional image with no diffusion sensitization (i.e., T2-weighted b0 images).

MEG recordings were acquired using an Elekta-Neuromag MEG system with 306 channels at 1000Hz sampling frequency and using an online band-pass filtered between 0.1 and 330Hz. The recordings consisted of 3 min resting state with eyes open (rEO) and 3 min resting state with eyes closed (rEC). During acquisition, participants remained seated inside the magnetically shielded room (VacuumSchmelze GmbH, Hanau, Germany). The head shape of the participants was acquired using a three-dimensional Fastrak digitizer (Polhemus, Colchester, Vermont), in addition to three fiducial points (nasion and left and right pre-auricular points) as landmarks. Four head position indicator (HPI) coils were placed on the participant’s scalp (two on the forehead and two on the mastoids) and their position was continuously monitored during the acquisition to allow for head position tracking. Last, two sets of bipolar electrodes were used to record eye-blinks and heart beats, respectively.

Informed consent was obtained prior to the recordings from all participants in accordance with the Declaration of Helsinki and the study received ethical approval from the ethics committee of the Universidad Complutense de Madrid.

### MEG processing

MEG recordings were preprocessed offline using the spatiotemporal signal space separation (tSSS) algorithm (Taulu and Hari, 2009), embedded in the Maxfilter software v2.2 (correlation limit of 0.9 and correlation window of 10 seconds), to eliminate magnetic noise and compensate for head movements during the recording. Continuous MEG data were preprocessed using the Fieldtrip toolbox (Oostenveld et al., 2011) in Matlab R2020b. An independent component analysis based on SOBI (Belouchrani, A., 1997) was applied to remove the eye-blink and cardiac signals from the data. Then, signals were visually inspected, and the remaining artefacts were identified and excluded from subsequent analysis.

Source reconstruction was performed using the Brainstorm toolbox (Tadel et al., 2011) in Matlab employing the minimum norm estimates (MNE) method (Dale and Sereno, 1993; Hämäläinen and Ilmoniemi, 1994) with *constrained dipoles* oriented normally to the cortical surface. This captures the typical orientation of the macrocolumns of pyramidal neurons (Tadel et al., 2019) and improves source localization accuracy (Dale and Sereno, 1993). Finally, source-space signals were averaged per region using the HCP parcellation scheme (Glasser et al., 2016) (see Supp. Table 1).

**Table 1.** JR-BNM parameters used in simulations.

| Parameter | Value | Unit | Description |
| --- | --- | --- | --- |
| $H_e$ | 3.25 | mV | Average excitatory synaptic gain |
| $H_i$ | 22 | mV | Average inhibitory synaptic gain |
| $\tau_e$ | 10 | ms | Time Constant of excitatory PSP |
| $\tau_i$ | 20 | ms | Time Constant of inhibitory PSP |
| $C_{pe}$ | 135 | | Average synaptic contacts: pyramidals to excitatory interneurons |
| $C_{ep}$ | 108 | | Average synaptic contacts: excitatory interneurons to pyramidals |
| $C_{pi}$ | 33.75 | | Average synaptic contacts: pyramidals to inhibitory interneurons |
| $C_{ip}$ | 33.75 | | Average synaptic contacts: inhibitory interneurons to pyramidals |
| $e_0$ | 0.0025 | $\text{ms}^{-1}$ | Half the maximum firing rate |
| $r$ | 0.56 | $\text{mV}^{-1}$ | Slope of the presynaptic function at $v_0$ |
| $v_0$ | 6 | mV | Potential when half the maximum firing rate is achieved |
| $p$ | variable | $\text{ms}^{-1}$ | Mean of random gaussian intrinsic noisy input |
| $\sigma$ | variable | $\text{ms}^{-1}$ | Standard deviation of random gaussian intrinsic noisy input |
| $g$ | variable | | Coupling factor for inter-regional communication |
| $s$ | 3.9 | m/s | Conduction speed for inter-regional communication |

### Alpha fluctuations

We used the MNE package (Gramfort et al., 2014) in Python 3.9 to extract a time-frequency representation (TFR) of each signal using Morlet wavelets (7 cycles) over a frequency range from 2 to 40 Hz in steps of 0.25 Hz. We computed the power spectrum by averaging the TFRs across time. The resulting spectra were modelled using the *fooof* package for Python (Donoghue et al., 2020) in the frequency range between 2 and 40 Hz. The model separates the periodic and aperiodic components of the spectrum and estimates the offset, exponent, and knee of the latter. From the periodic components identified, we selected that with the highest power in alpha band (8 - 12 Hz) as the individual alpha frequency (IAF). In the case of no alpha peak detected by fooof, IAF was considered to be 10Hz.

To evaluate the fluctuations of alpha power over time, we selected a frequency band of the TFR at IAF +/- 2 Hz. The TFR values were then averaged in band, and the results were scaled by a factor of 1e-19. Finally, we evaluated the distribution of power values by computing a histogram of 200 bins following Freyer et al. (2009). We evaluated the fit of two types of exponential functions to the distribution of powers: unimodal exponential *P* (*x*) = *λe^−λx^*, and bimodal exponential *P* (*x*) = *wλ*_1_*e^−λ^*^1^ *^x^* + (1 − *w*)*λ*_2_*e^−λ^*^2^ *^x^*. The goodness of fit was evaluated using the Bayesian Information Criteria (BIC): *BIC* = *k* ln(*n*) − 2 ln(*L*), where k is the number of parameters used (1 for unimodal and 3 for bimodal), n is the number of datapoints, and L represents the likelihood of the model.

### Functional connectivity

The FC was estimated with two different metrics: the corrected imaginary part of the phase locking value (ciPLV; (Bruña et al., 2018)) for phase synchrony, and the corrected version of the amplitude envelope correlation with pairwise signals orthogonalization (cAEC; (Hipp et al., 2012; O’Neill et al., 2015)) for amplitude synchrony. Both measures correct for source leakage and volume conduction.

To compute these metrics, we epoched the signals using a sliding window approach with windows of 1 second -and additional padding of 1 second at each extreme of the window- and an overlap of 0.5 seconds. We filtered the epoched data in 4 different frequency bands: theta (4-8 Hz), alpha (8-12 Hz), beta (12-30 Hz) and gamma (30-45 Hz); and computed the Hilbert transform of the signals. The differences in the imaginary part of the Hilbert’s phases were used to compute ciPLV, while the orthogonalized envelopes of that transformation were correlated to get the cAEC. This process was performed per window, subject, and condition for each pair of regions in the DMN (see Supp. Table 2).

### Structural connectivity

DWI data was processed using DSI Studio (http://dsi-studio.labsolver.org). Firstly, the DWI data was rotated to align with the AC-PC line. The restricted diffusion was quantified using restricted diffusion imaging (Yeh et al., 2016). The diffusion data were reconstructed using generalized q-sampling imaging (Yeh et al., 2010) with a diffusion sampling length ratio of 1.25. The tensor metrics were calculated using DWI with a b-value lower than 1750 s/mm². A deterministic fiber tracking algorithm (Yeh et al., 2013) was used with augmented tracking strategies (Yeh, 2020) to improve reproducibility, using the whole brain as seeding region. The anisotropy threshold was randomly selected. The angular threshold was randomly selected from 15 degrees to 90 degrees. The step size was randomly selected from 0.5 voxels to 1.5 voxels. Tracks with lengths shorter than 10 or longer than 300 mm were discarded. A total of 1 million seeds were placed.

The HCPex v1.1 atlas (Glasser et al., 2016; Huang et al., 2021) was used as the volume parcellation atlas that complements the HCP atlas by including subcortical structures (see Supp. Table 1). Finally, two structural connectivity (SC) matrices were constructed according to the count and average length of the streamlines connecting two regions.

### Brain network model

SC matrices served as the skeleton for the BNMs implemented in The Virtual Brain (Sanz-Leon et al., 2015) where regional signals were simulated using JR NMMs (Jansen and Rit, 1995). The JR is a biologically inspired model of a cortical column capable of reproducing alpha oscillations through a system of second-order differential equations (see Table 1 for a description of parameters):

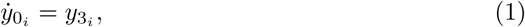

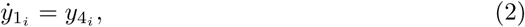

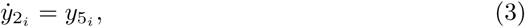

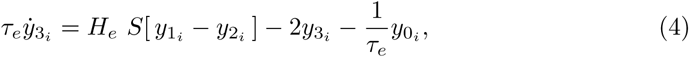

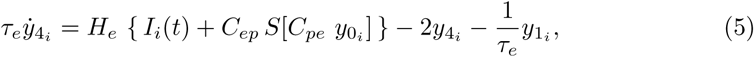

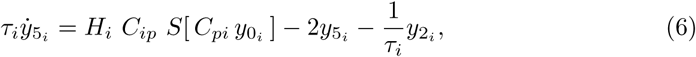

where

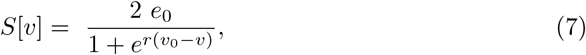

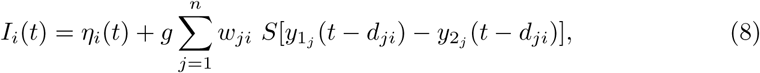

for *i* = 1*, …, N*, where *N* is the number of simulated regions. The inter-regional communication introduces heterogeneity in terms of connection strength *w_ji_* and conduction delays *d_ji_* between nodes i and j where: 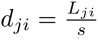, with *L_ji_* being the length of the tract from node i to node j, and *s* representing the (global) conduction speed.

This model represents the electrophysiological activity (in voltage) of three subpopulations of neurons: pyramidal neurons (*y*_0_), excitatory interneurons (*y*_1_), and inhibitory interneurons (*y*_2_) and their derivatives (*y*_3_, *y*_4_, and *y*_5_, respectively). These subpopulations are interconnected through an average number of synaptic contacts (*C_ep_, C_ip_, C_pe_* and *C_pi_*), and integrate external inputs from other cortical columns. The intra- and interregional communication is mediated through firing rates, which are determined by a sigmoidal function *S* converting voltage inputs into firing rates (eq. 7). The shape of the sigmoidal function is determined by its steepness *r*, its half-maximum *e*_0_, and the voltage required to reach half-maximum *v*_0_. The postsynaptic potential amplitudes (H_e_, H_i_), and time constants (*τ_e_, τ_i_*) shape the oscillatory behaviour of subpopulations’ voltages.

The input (*I_i_*(*t*)) represents two main drivers of activity in the NMMs: inter-regional communication and an intrinsic noisy input. The global coupling *g* scales the weight of the tracts connecting the brain regions of SC, as shown in (8). The intrinsic noisy input is defined as a local and independent Gaussian noise *η_i_*(*t*) ∼ N (*p, σ*).

### Simulations

All simulations were 60 seconds in length, from which we discarded 8 seconds to avoid initial transients. Sampling frequency was 1 kHz. Two types of simulations were carried out: single node simulations and network simulations. For the latter, we downsampled the SC by averaging the 426 areas of the HCPex atlas into 66 regions defined in Supp. Table 1.

To assess whether the simulated signals followed an exponential distribution, we computed the log(likelihood) of the simulated data given the exponential function derived from empirical recordings. To do this, we first normalized the empirical time-frequency representations (TFR(*α*)) within each region and then recomputed the parameters of the corresponding exponential distributions. This normalization ensured that both simulated and empirical TFR(*α*) were directly comparable on the same scale, allowing for a robust evaluation of their distributional properties.

### Statistics

To assess statistical differences between conditions and brain regions in the parameterization of the exponential functions derived from empirical data, we conducted a series of Friedman’s tests as a robust alternative to one-way repeated-measures ANOVAs, given that homocedasticity assumption was not met.

Additionally, we performed a set of Spearman’s correlation tests to relate the spectral characteristics of the signals (i.e., IAF, power, and aperiodic exponent) to the difference in BIC between unimodal and bimodal exponential functions, and to the value of BIC derived from the best performing model.

Finally, we evaluated the relationship between the FC and the fluctuations of alpha power by means of Spearman’s correlations. To this end, we downsampled in time the TFR(*α*) normalized per subject to match the time resolution of the FC analysis (windows of 1 second with steps of 0.5). This process generates a one dimensional array per region, which represents the level of alpha power across time. Then, for each pair of regions in the DMN, their average TFR(*α*) was correlated with the FC of the connection in time for each frequency band. Once with a TFR(*α*)-FC(band) correlation per metric and band, we averaged the correlations values per subject and performed an independent samples t-test against a normal distribution centred at zero.

Corrections for multiple comparisons were performed with Bonferroni method.

## Results

### The distribution of alpha power fluctuations follows an exponential function

To evaluate alpha power fluctuations over time, we performed a time-frequency analysis of the signals, filtering them in the IAF +/- 2 Hz range, and averaging out frequencies to obtain an array of power values over time (see Fig. 1B). Then, we calculated the distribution of those power values by elaborating a histogram with 200 equally sized bins following Freyer et al. (2009). The resulting distributions followed exponential functions that in some cases were better represented by bimodal exponentials (see Fig. 1A - green regions in BIC). An example of these bimodalities, can be appreciated in the axis transformations shown in Figure 1B, where the distribution of power for rEC was better captured by a bimodal exponential function [BIC = 432534] than by its unimodal counterpart [BIC = 474451].

**Figure 1.**
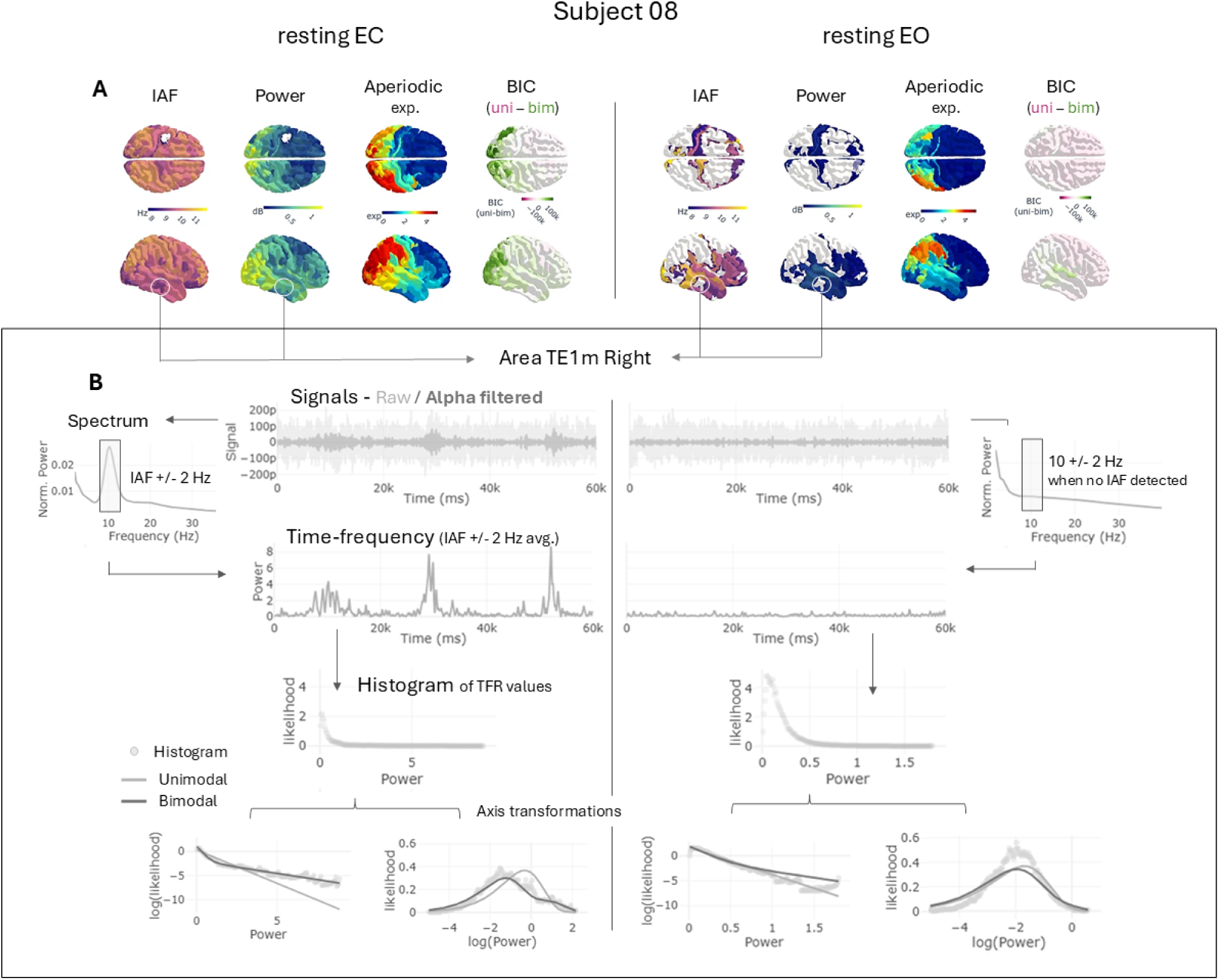
Pipeline of analysis for a sample subject (sub-08) in both rEC (left) and rEO (right) conditions. A) Results of the spectral modelling including IAF, power, and the exponent of the aperiodic component. White regions in IAF and power represent non-detected alpha peaks. In these cases, we analysed the 10 +/- 2 Hz frequency band. BIC column shows the difference in goodness of fit between the unimodal and bimodal exponential functions (positive values in green favour bimodality). B) Alpha fluctuations analysis for one sample region (TEm1 R) both in rEC and rEO. First row, showing the signals (raw and filtered in alpha band); second, the spectra derived from the raw signal that is used to detect the IAF; third, TFR analysis in the IAF +/- 2 Hz frequency band averaged through frequencies to obtain a single array of alpha power in time; fourth, histogram with 200 equally sized bins using the data array of alpha power (each dot represents a bin); fifth, axis transformations of original exponential distributions helps to evaluate visually the presence of bimodalities. The fit of the unimodal and bimodal functions are shown in light and dark gray lines, respectively. Shared y-axis for rEC and rEO.

In addition to that procedure, we evaluated the average spectral characteristics of the signals by computing their frequency spectra, modelling the aperiodic component, and measuring alpha frequency peak and power (Fig.1A). Note that some regions do not show a significant alpha peak over the aperiodic component as modelled by fooof, specially in rEO (see Fig. 1A - blank regions in IAF and power). In those cases, the IAF used for the analysis was 10 +/- 2 Hz. Interestingly, the fluctuations of alpha in these regimes could also be accurately approximated by exponential distributions (see Fig. 1B - rEO).

### Bimodal distributions are associated with high alpha power in posterior regions

Looking into the group averaged results, we observed that the bimodal distributions appeared most frequently over posterior regions, both in rEC and rEO (see Fig. 2A). We evaluated statistically this observation by performing two Friedman’s tests on the BIC differences between the unimodal and bimodal exponential fits using condition and space (i.e., brain region) as factors. We found significant differences for space [*χ*^2^(359, 42)=817.42, p-corr*<*0.0001, W=0.455] but not between conditions [*χ*^2^(1, 42)=0.857, p-corr=1, W=0.02]. Bimodalities appeared most often in posterior regions, but they were not always present, 11.90% of subjects showed no bimodality in rEC and 2.39% in rEO. In average, 15.42% (std 16.29%) of all brain regions were better modeled with bimodal exponentials. Frontal regions, in contrast, showed a clear tendency towards unimodality (see Fig. 2A).

**Figure 2.**
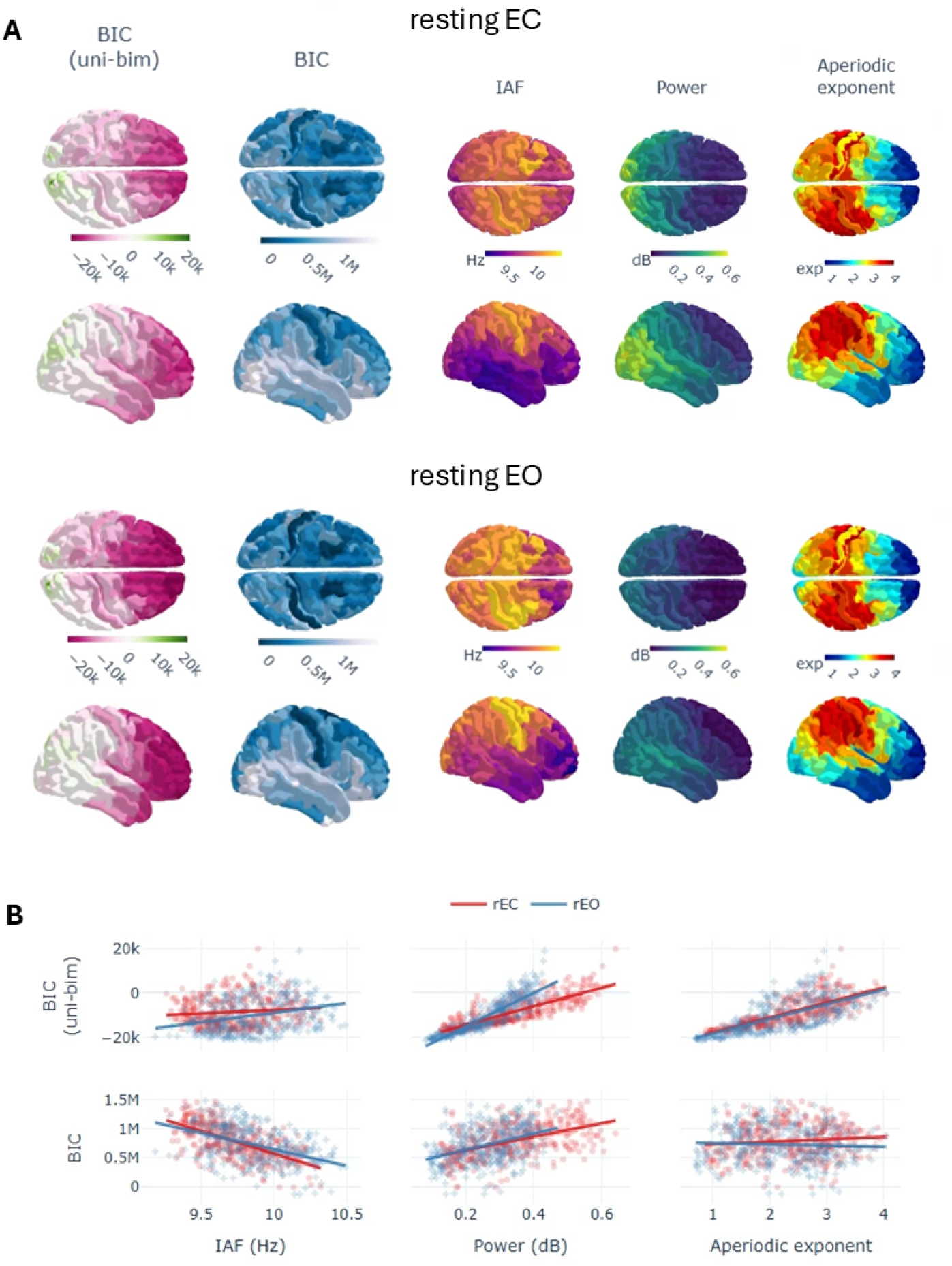
Regional characterization of the exponential modelling and spectral variables in rEC/rEO. Values were averaged through subjects. A) Topological descriptions including the BIC difference between the unimodal and bimodal exponential fits (positive values in green favouring bimodality), the BIC value of the best fitted model per region (lower values in dark blue indicate better fit), and three spectral variables including IAF, alpha power and aperiodic exponent. B) Relationship between the exponential modelling variables and spectral ones. Note again that positive values in the BIC(uni-bim) favour bimodality and that lower BIC values imply better model fit.

Furthermore, we found strong correlations for the tendency towards bimodality and the spectral characteristics of the signal, specially for alpha power both in rEC [r(360)=0.852, p-corr*<*0.0001] and in rEO [r(360)=0.923, p-corr*<*0.0001] (see Fig. 2B - first row) but also for the aperiodic exponent in rEC [r(360)=0.82, p-corr*<*0.0001] and rEO [r(360)=0.71, p-corr*<*0.0001]. A slightly different picture was derived from the relationships between the spectral characteristics and the goodness of fit of the (best performing)exponential model. In this case, we observed inverse correlations for IAF [rEC: r(360)=-0.63, p-corr*<*0.0001; rEO: r(360)=-0.48, p-corr*<*0.0001] and direct moderate correlations for power [rEC: r(360)=0.53, p-corr*<*0.0001; rEO: r(360)=0.4, p-corr*<*0.0001] (see Fig. 2B - last row). Correlations for the aperiodic exponent did not result significant. Interestingly, when comparing directly spectral characteristics between rEC and rEO, we only found a significant difference between conditions for power [*χ*^2^(1, 42)=13.71, p-corr*<*0.002, W=0.326] with higher alpha for rEC, but not for aperiodic exponent [*χ*^2^(1, 42)=0.095, p-corr=1, W=0.0022] and IAF [*χ*^2^(1, 42)=1.523, p-corr=1, W=0.036] (see Fig 2.1).

In summary, these results show that higher alpha power as measured in the spectrum is related to bimodalities in the distribution of alpha fluctuations. Also, that the higher the power the more deviated it is the distribution from an exponential function, given by the decrease in goodness of fit. Finally, that regions with higher spectral alpha frequencies are associated to distributions closer to the exponential.

### The FC of the DMN is directly related to the amplitude of alpha oscillations

In this section, we studied how the fluctuations of alpha power are related to the activity of the DMN. We expected to find differences in network activation depending on the levels of alpha, and more specifically, to find a negative relationship between the power of alpha and the FC in the DMN. We evaluated the FC within this network using two complementary measures: ciPLV to evaluate phase synchrony and cAEC to evaluate amplitude synchrony. Then, we correlated FC values with the alpha power in time for each pair of DMN regions. Finally, we averaged the correlation values per subject within the network, and made a group test against zero to evaluate for significant relationships between the variables.

Both metrics yielded complementary insights on the relationship between alpha power fluctuations and FC (Fig. 3). With ciPLV, we observed a prominent direct relationship between alpha power and FC in alpha band [rEC: T(39)=6.95, p-corr*<*0.001, Cohen’s d=1.09; rEO: T(39)=6.06, p-corr*<*0.001, Cohen’s d=0.96] and beta [rEC: T(39)=5.72, p-corr*<*0.001, Cohen’s d=0.90; rEO: T(39)=5.86, p-corr*<*0.001, Cohen’s d=0.92] bands. Interestingly, for FC in theta band we found a tendency towards negative relationships that resulted significant for rEC [T(39)=-4.12, p-corr*<*0.005, Cohen’s d=0.65]. With cAEC, FC resulted in a direct and significant relationship for alpha FC in rEO [T(39)=5.60, p-corr*<*0.001, Cohen’s d=0.88] but not in rEC[T(39)=2.24, p-corr=0.48]. In this case, FC in theta band showed direct and significant relationships [rEC: T(39)=4.70, p-corr*<*0.001, Cohen’s d=0.74; rEO: T(39)=4.30, p-corr*<*0.005, Cohen’s d=0.68] while beta and gamma did not.

**Figure 3.**
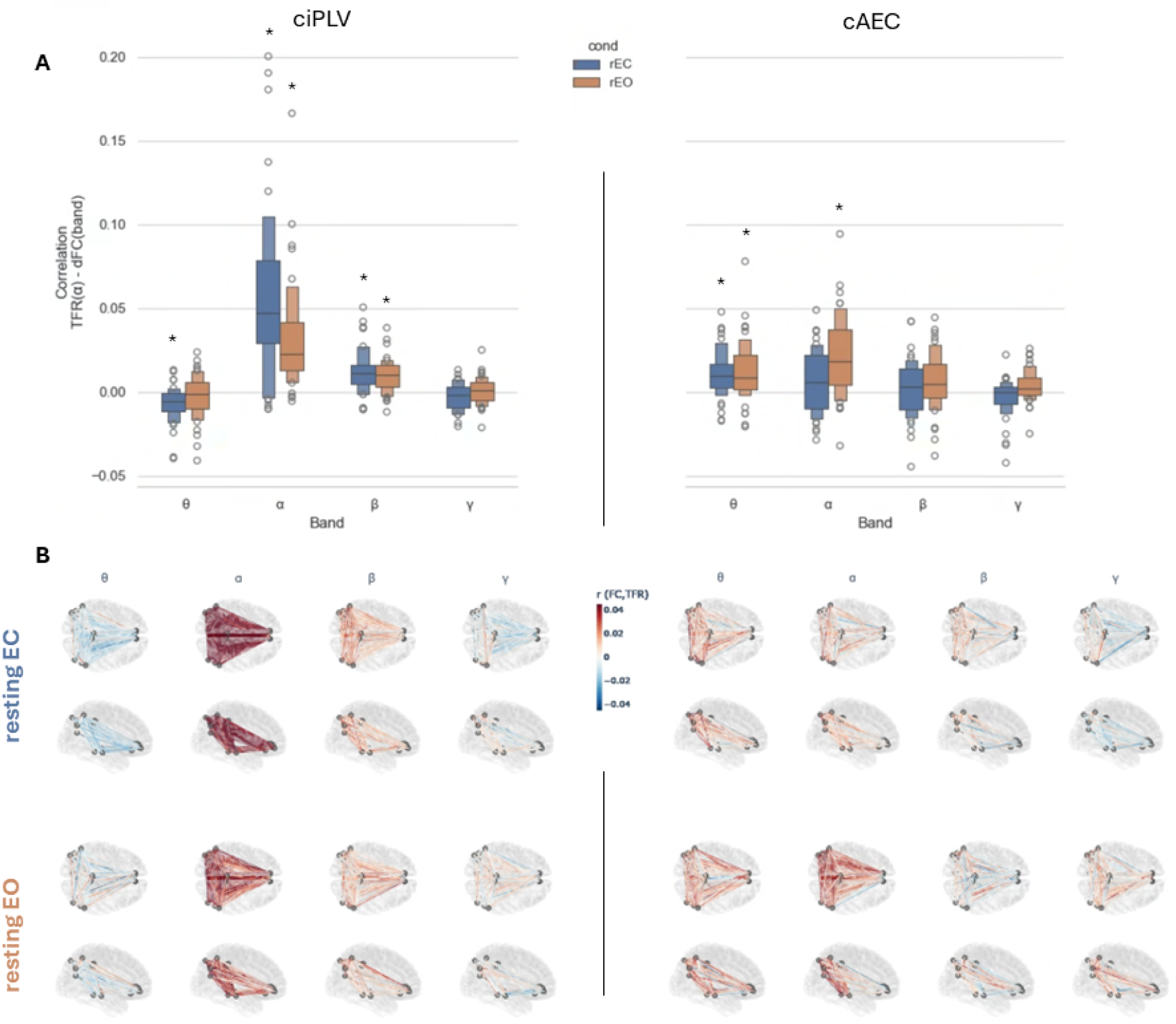
A) Correlations between alpha power and FC in the DMN per band averaging out specific connections (each datapoint represent one subject), differentiating by FC metric (ciPLV/cAEC) and condition (rEC in blue and rEO in orange). B) Correlations between alpha power and FC per band averaging out subjects to show each connection in the DMN. In color, the direction of the correlation with positive in red, negative in blue. (*) corresponds to statistical significance after correction for multiple comparisons, with p-corr<0.01.

### The fixed points in Jansen-Rit reproduce better alpha power fluctuations

The JR model is widely used to simulate electrophysiological alpha oscillations. Here, we evaluate to what extent and under which parameterizations the JR reproduces the empirically observed power fluctuations. We followed a constructive approach by simulating first a single node and then whole brain networks to explore the parameter spaces and model performance.

The results of single node simulations showed better fits to the average empirical exponential function when the model was operating in a regime of attracting fixed points (steady states) (see Fig. 4 - 5th column) both before the saddle node (parameter *p<*0.11) and after the supercritical (parameter *p >* 0.33) bifurcations of the JR. Between these two critical points, the JR exhibits limit cycle behaviours, with a worse fit to the empirical exponential. Also, increasing the levels of noise (i.e., *σ*) in the limit cycle regimes increased the goodness of fit.

**Figure 4.**
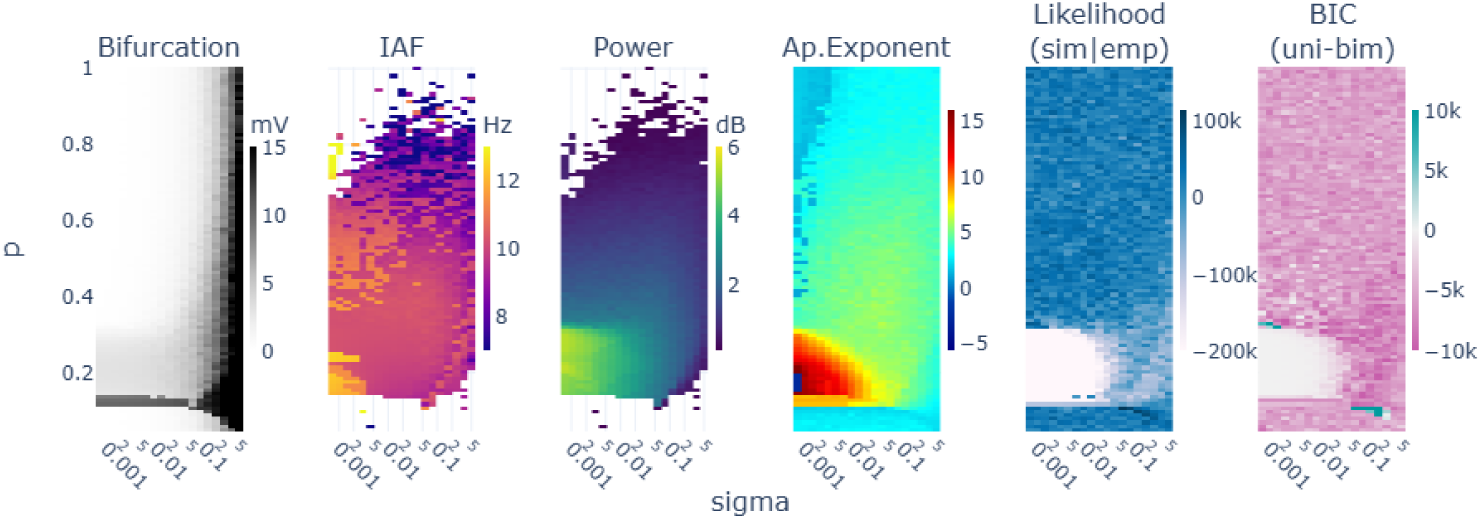
Parameter space explorations for a single node varying the mean intrinsic input (p) and standard deviation (sigma). In columns, 1) the bifurcation as the signal’s max-min voltage, 2) the alpha frequency peak, 3) the peak’s power and 4) aperiodic exponent as modelled with fooof toolbox, 5) the log(likelihood) of the unimodal exponential fit, and 6) the BIC difference between the unimodal and bimodal exponentials -the higher favours unimodal distributions-. Blank values in columns 2) and 3) correspond to undetected alpha peaks by fooof modeling.

We extracted a subset of samples from the parameter spaces to get a better grasp of the simulation results. In Fig. 5, we show the signals, TFR, averaged spectrum and exponential modelling results for three different parameter combinations (i.e., sigma=0.01, p=[0.09, 0.22, 0.44]). The simulation performed within the regime of stable limit cycle (e.g. p=0.22) resulted in less biologically plausible alpha rhythm (Likelihood(sim—emp)=-289256.28) with a constant high power and little fluctuations, compared to the regimes with stable fixed points (Likelihood(sim—emp)=[38316.90, 22853.69]; Fig. 5 - see the bifurcation diagram as reference).

**Figure 5.**
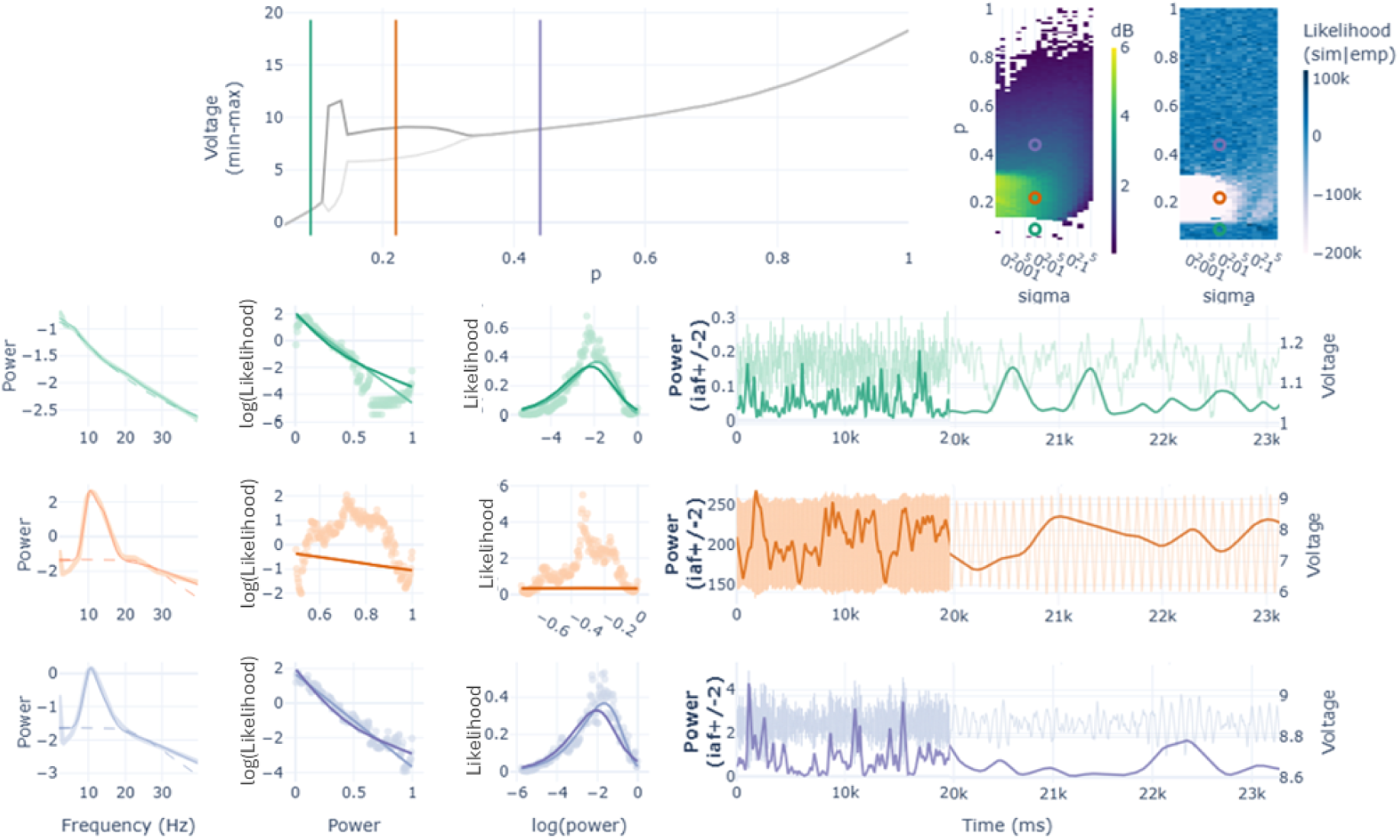
Three samples of the single node simulations with p=[0.09, 0.22, 0.44] and sigma=0.01. First row shows the bifurcation and two parameter spaces as reference for the picked simulations (in coloured vertical lines and open circles). The following rows show: 1) the power spectrum calculated from the simulated signals (thick line), the modelled spectra (thin line), and the modelled aperiodic component (dashed line); 2) the exponential function in two different coordinate spaces (i.e., Linear-Log and Log-Linear) with the scatter representing the histogram of the simulated TFR(α) and the lines representing both the unimodal (light colour) and the bimodal exponential models (dark colour); 3) the raw signal (light colour) and the TFR(α) values (dark colour) in two different timeframes: 0-20 seconds for a broader overview, and 20-23 seconds for finer detail.

Regarding the fixed point states, we noticed that the fixed point regimes after the supercritical bifurcation (parameter *p >* 0.33) show an alpha peak over the aperiodic component (see Fig. 5 - first column, last row) in contrast to the fixed point regime before the first saddle node bifurcation in which an alpha peak does not appear (parameter *p<*0.11; see Fig. 5 - first column, green trace).

Regarding bimodalities, we expected them to appear at the critical points where the system could theoretically transition between different states of alpha. In this line, we found two sets of simulations favouring bimodal exponentials: one related to the first saddle node bifurcation (p≈0.1025) and another related to the supercritical one (p≈0.33). At the supercritical bifurcation with low noise (Fig. 4 - last column [p≈0.33, sigma≈0.005], in blue) we found a bimodal behaviour that was artificially produced by the slowly decaying initial transient that overcome the discarded initial 8 seconds of simulation (see Fig 5.1 and Fig 5.2). In contrast, at the saddle node bifurcation with higher noise (Fig. 4 - last column [p≈0.1025, sigma≈0.1]) an actual switching behaviour was found between the fixed point and the limit cycle regimes of the JR (see Fig 5.3). The switching consisted in events/periods of high voltage alpha fluctuations intertwined with periods of a low voltage noisy signal characteristic of the fixed points.

These results suggest that the fixed point regimes reproduce the empirically-observed alpha power fluctuations, and that the model do not show biologically plausible bimodalities in the explored parameter space.

### Theory and simulations favour post- supercritical fixed points to reproduce electrophysiological alpha oscillations

We simulated the whole brain network to evaluate how the interactions between nodes could affect the fluctuation dynamics. We fixed the noise level (i.e., sigma=0.001) and explored both the coupling factor (i.e., g) and the mean intrinsic input (i.e., p) parameters. The simulations were performed using the averaged SC of the sample of subjects.

In the resulting parameter spaces, we found a bifurcation with a diagonal shape (see Fig. 6), what is not surprising given the roles of *g* and *p* as bifurcation parameters in the model. Apart from this difference, the spectral results and the exponential fits followed similar lines of single node simulations. The worst fit of the unimodal exponential was found within the limit cycle of the JR. Before and after the limit cycle, the fluctuations of alpha power adjusted better to an exponential function. Although higher rates of bimodality might have been expected in these network simulations due to node interactions, they were in fact rare and sparsely distributed across the two main bifurcations of the model. In general, the model generated alpha fluctuations that were better captured by unimodal exponential distributions.

**Figure 6.**
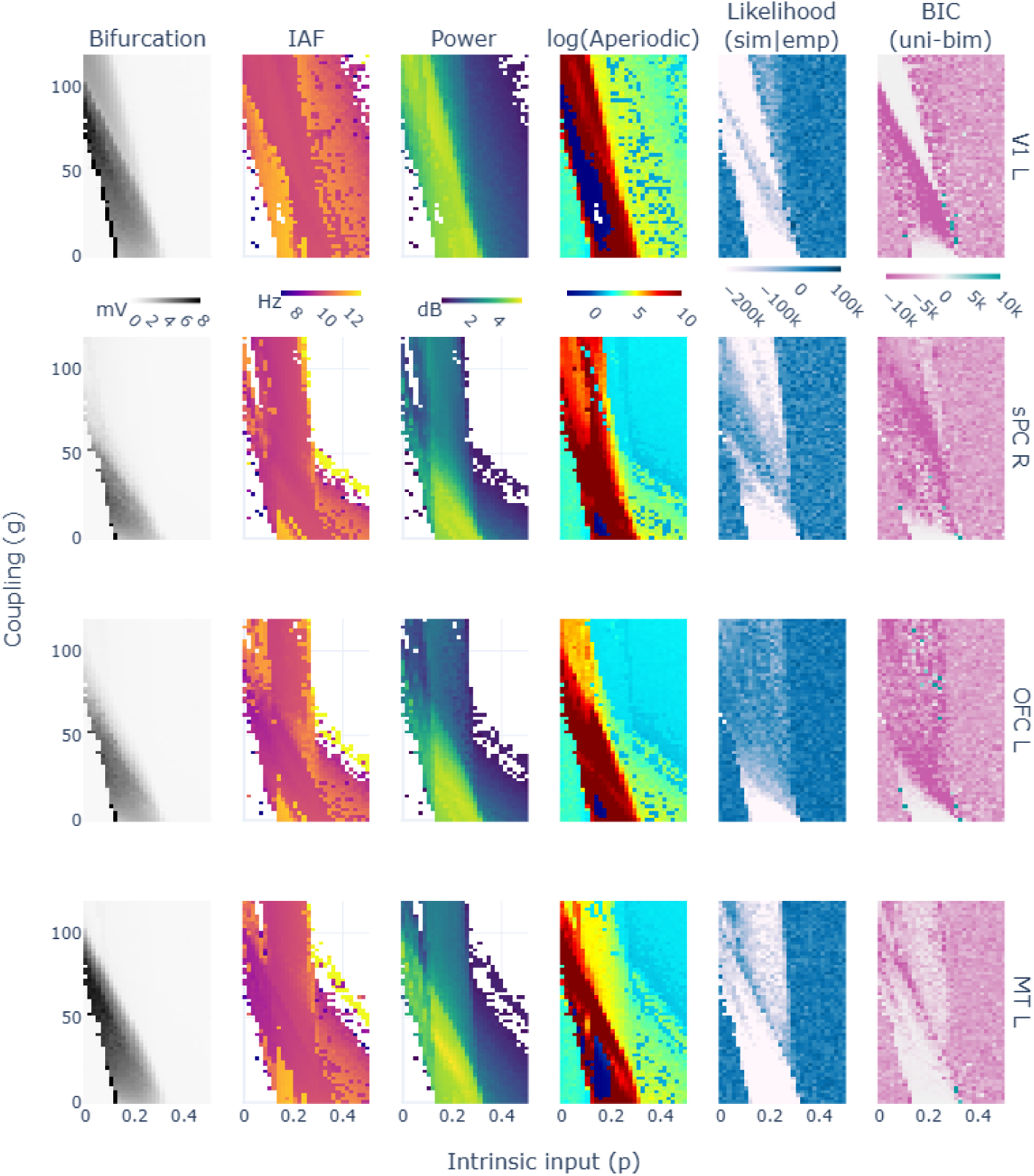
Parameter space explorations for network simulations with a fixed sigma=0.001 and varying p and g. In columns, 1) the bifurcation as the signal’s max-min voltage, 2) the alpha frequency peak, 3) the peak’s power and 4) aperiodic exponent as modelled with fooof toolbox, 5) the BIC of the unimodal exponential fit, and 6) the BIC difference between the unimodal and bimodal exponentials -the higher favours unimodal distributions-. Not shown values in columns 2 and 3 correspond to undetected alpha peaks by fooof modeling. Each row corresponds to analysis from simulated signals in different brain regions. Abbreviations: Orbito-Frontal Cortex (OFC), Medial Temporal Cortex (MT), Superior Parietal Cortex (sPC), Primary Visual Cortex (V1).

Similarly to single node analysis, alpha peaks were found in post- supercritical bifurcation fixed points, while no alpha peak was detected in the pre- saddle node ones (see 6 - Power column; and Fig 6.1). Also, the spectral alpha power tends to decay as the node progresses further into the post- supercritical bifurcation regimes.

These results suggest that the fixed points in the post-supercritical bifurcation reproduce better electrophysiological alpha fluctuations than the typically used limit cycle operation state. We argue this given that: 1) fixed points reproduce better an exponential function of alpha fluctuations than the limit cycles, 2) the pre- saddle node fixed points do not show alpha activity in contrast to post- supercritical fixed points, and 3) the progressive reduction in alpha power through the post- supercritical bifurcation matches the theory of alpha claiming that it reflects inhibition / an idling state, in which case, the increase of input to the region would generate a reduction in alpha power.

## Discussion

In this study, we have characterized the temporal fluctuations of alpha power, modelling them as exponential distributions and examining topological differences between rEC and rEO conditions. Additionally, we investigated the relationship between alpha power fluctuations and FC within the DMN, and assessed the performance of the JR neural mass model to accurately reproduce the shape of the empirically observed fluctuations, both in isolation and as part of a broader brain network model.

Our findings corroborate prior research suggesting exponential distributions of alpha power fluctuations (Freyer et al., 2009), with bimodal patterns linked to high-power states in posterior regions. In our sample, these bimodalities appeared less frequent and less pronounced than previously reported (Freyer et al., 2009). In their work, Freyer et al. (2011, 2009) proposed a mechanistic explanation for this phenomenon, suggesting that the thalamocortical system operates in a subcritical Hopf bifurcation regime that allows it to switch between two states of alpha -low and high alpha amplitude-. Besides this hypothesis, we believe that those two states of alpha could reflect functionally different modes of the resting state. Resting state is a complex and dynamic condition in which participants are instructed to remain seated and relaxed, without engaging in any specific task. However, spontaneous mental activities, such as mind-wandering, episodic memory recall, and internal thought processing may occur involuntarily during these periods (Raichle et al., 2001; Smallwood and Schooler, 2015; Christoff et al., 2016). The switching over time been an internal focus of attention -as in the case of mind wandering- and an external one might contribute to the differentiation of two modes of alpha (Fox et al., 2015; Sadaghiani and Kleinschmidt, 2016). Finally, also from the technical point of view, we believe that volume conduction and source leakage could have intensified the observed bimodal effects by mixing signals from different sources. The use of MEG data in this study with higher spatial resolution and the use of source reconstruction methods might have reduced the prominence of this phenomenon.

Regarding the relationship between alpha power fluctuations and dynamical FC, both phase (ciPLV) and amplitude (cAEC) metrics of FC showed direct relationships *within* the DMN. For this matter, *where* each aspect is evaluated becomes key. For instance, some studies have related the activation of the DMN with alpha power *in the visual cortex*, showing also direct relationships (Kim et al., 2023; Clancy et al., 2021; Mo et al., 2013). In our case, we have evaluated both alpha power and FC *within* the regions of the DMN. That’s why, we hypothesized to find an inverse relationship (Scheeringa et al., 2012; Bowman et al., 2017), based on multimodal fMRI-EEG studies showing that higher alpha power is associated to lower BOLD signal (Goldman et al., 2002; Laufs et al., 2003; Pang and Robinson, 2018). Only theta band ciPLV in rEC resulted in significant negative relationships. These findings raise important questions regarding the intricate relationship between different neuroimaging modalities and their respective FC measures, emphasizing the need for integrative approaches to bridge gaps between electrophysiological and hemodynamic signals (Logothetis, 2008; Ritter and Villringer, 2006; Mulert, 2013). If high alpha power —associated with reduced BOLD signal— suggests a state of inhibition or idling (Jensen, 2024; Pfurtscheller et al., 1996), in the DMN it would suggest a state of disengagement from mind-wandering. How could electrophysiological measures of FC within that network concurrently indicate an increase? What would FC mean in that context? Future research should further investigate the mechanisms underlying these interactions and their implications for cognitive and neural processing.

On the computational side of this work, our results suggest the use of the JR’s fixed points -instead of its limit cycles or critical points- to simulate electrophysiological dynamics. Furthermore, we found a better performance for the post- supercritical fixed points over the pre- saddle node ones. This is because (1) fixed points reproduced better the exponential distributions of alpha fluctuations, (2) post- supercritical fixed points showed alpha power over the aperiodic component of the spectrum -as expected in resting-state- unlike pre- saddle node ones, and (3) the relationship between the increasing input to a node and the reduction in alpha power aligns with experimental data from fMRI-EEG co-registers reporting anti-correlations between alpha power and neuronal metabolism (Goldman et al., 2002; Laufs et al., 2003; Pang and Robinson, 2018). Across this range of fixed points, different levels of alpha power can be simulated. As the bifurcation parameter increases, the average alpha power decreases, potentially reaching a regime where no distinct alpha peak emerges above the aperiodic component.

It has been argued that the neural mass models used in whole-brain modeling might be parameterized at criticality, associating the critical points of a dynamical system with the criticality described in empirical neurophysiological data (Breakspear, 2017; Deco and Jirsa, 2012). At the critical point, the system would be able to switch between different modes of operation, providing flexibility and adaptability to the system. This idea has some support in the context of electrophysiological studies (Fuscà et al., 2023; Fontenele et al., 2019; Shew and Plenz, 2012), although still under discussion (Destexhe and Touboul, 2021). In either case, the extent to which the criticality in electrophysiology is well represented by the critical points of a dynamical system remains to be explored. In our data, we found two groups of single-node simulations showing bimodalities, and interestingly, both were parameterized at critical points of the system. However, in none of them the bimodality was due to biologically plausible transitions between modes of operation. In one of the groups, the bimodality was explained by the decaying initial transients of the system while, in the other, nodes were randomly switching between the pre- saddle fixed points and the limit cycle behaviour of the JR, resulting in signals that do not resemble M/EEG recordings. We believe that a further understanding of criticality in electrophysiology and dynamical system is needed to better align simulated dynamics to empirical data.

This work advances the understanding of alpha brain activity by elucidating its dynamic properties and their implications for neural processing. Furthermore, it establishes a robust framework for assessing the biological plausibility of simulated neural dynamics, offering a systematic approach to bridging computational models with empirical data. By integrating theoretical insights with empirical validation, this framework enhances the reliability of simulations as tools for investigating brain function and dysfunction, while also offering new avenues to understand alpha rhythms and its involvement in cognition.

## Supporting Information

**Fig 2.1.**
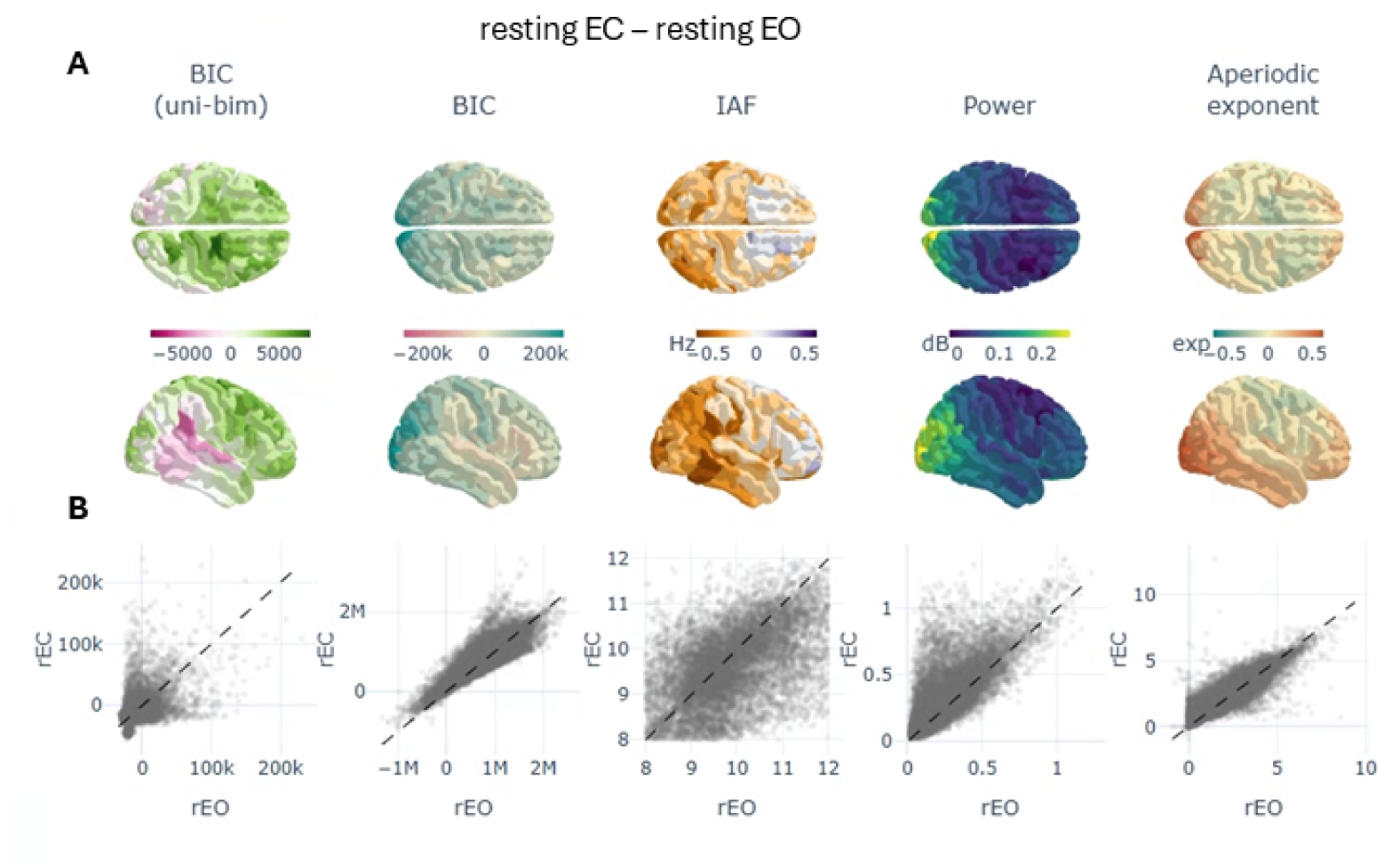
Intra-subject differences between rEC and rEO in the exponential modelling results and spectral variables. Values were averaged through subjects. A) Topological descriptions including the BIC difference between the unimodal and bimodal exponential fits (positive values in green indicate a tendency in rEC towards bimodality); the BIC value of the best fitted model per region (blue regions indicate a better fit in rEO than in rEC); and three spectral variables including IAF, alpha power and aperiodic exponent. B) Scatter plots with a dot per subject and region. The dashed line as 1:1 reference. Dots over the reference line represent increases in rEC.

**Fig 5.1.**
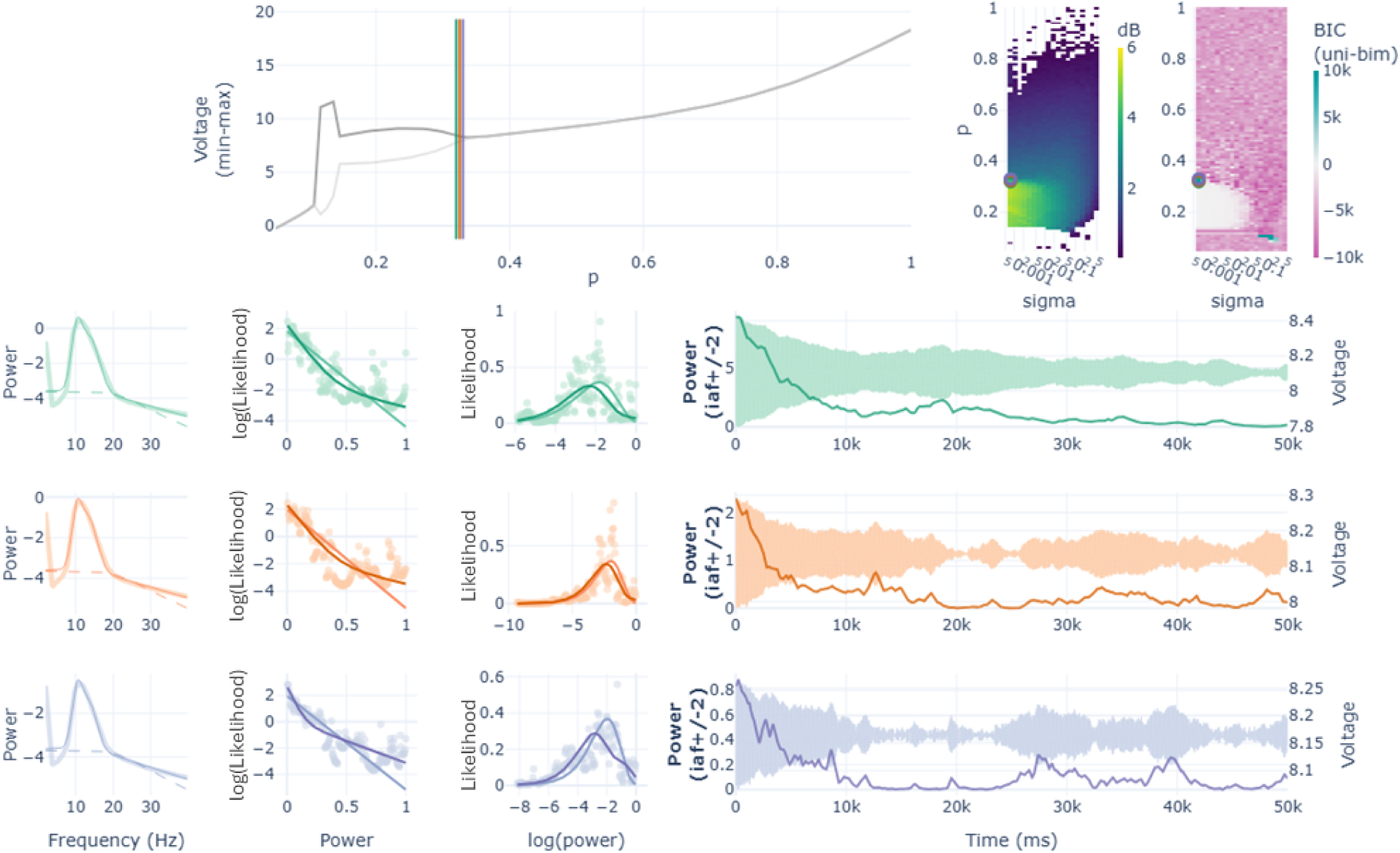
Samples of bimodalities found in single node simulations (Fig. 4) at the critical point of the subcritical bifurcation (p=[0.32, 0.325, 0.33]) with low noise (sigma = 0.00075). All the traces show a decaying initial transient that overcame the discarded initial 8 seconds of simulation. All the samples fitted better bimodal exponentials: green trace [BIC: unimodal = −83127.95; bimodal = −84328.11], orange trace [BIC: unimodal = −98014.40; bimodal = −99485.34], and blue trace [BIC: unimodal = −96800.84; bimodal =-100683.70]. Note how the initial transient becomes less evident as the model gets into the post-subcritical fixed point state.

**Fig 5.2.**
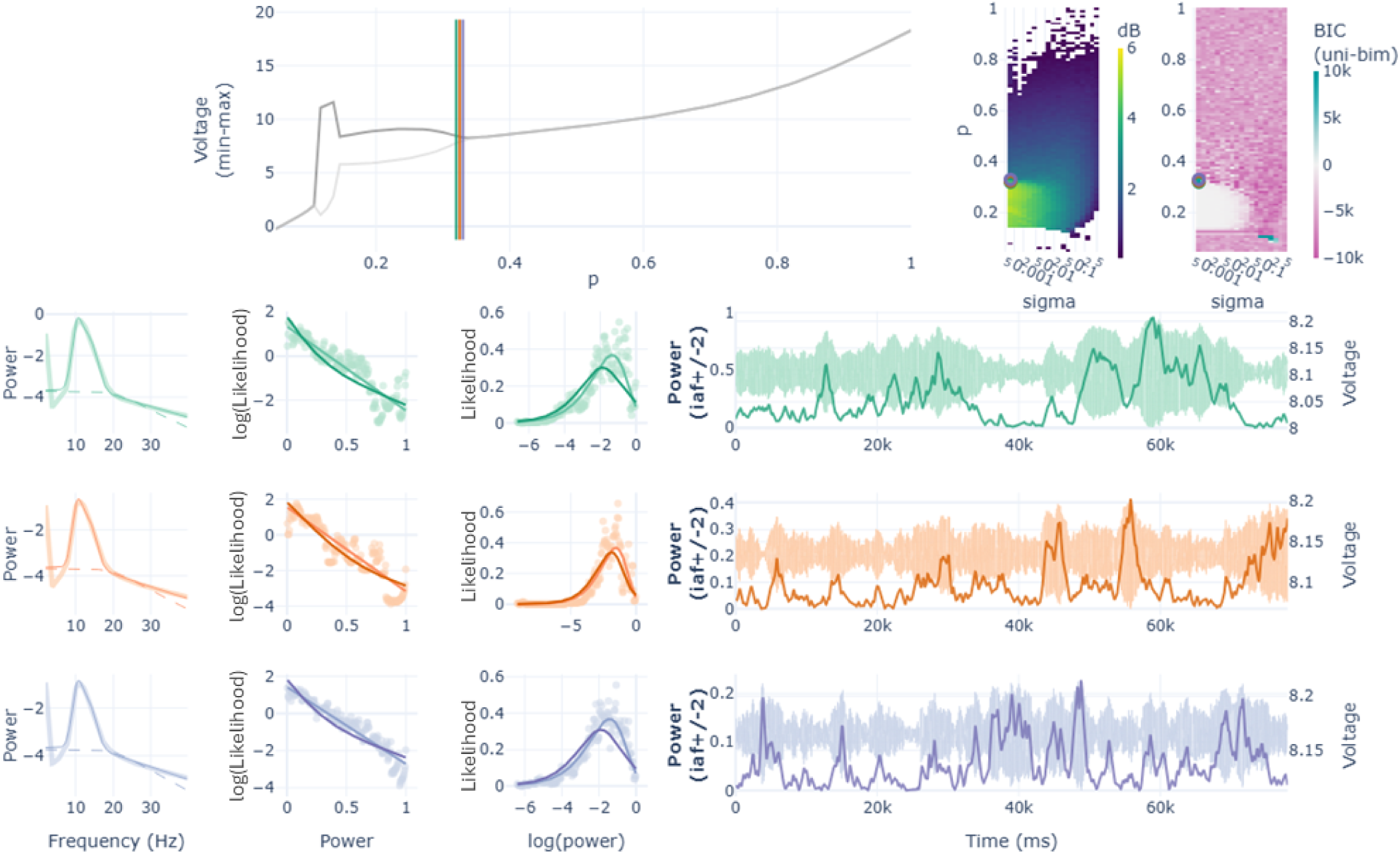
Same as in Fig 5.1 discarding longer transients (40 seconds): single node simulations at the critical point of the subcritical bifurcation (p=[0.32, 0.325, 0.33]) with low noise (sigma = 0.00075). With transients discarded bimodalities appeared less frequently. Here, all the samples fitted better unimodal exponentials: green trace [BIC: unimodal = −52965.25; bimodal = −39361.56], orange trace [BIC: unimodal = −86116.75; bimodal = -77302.37], and blue trace [BIC: unimodal = −65850.80; bimodal = −53257.19].

**Fig 5.3.**
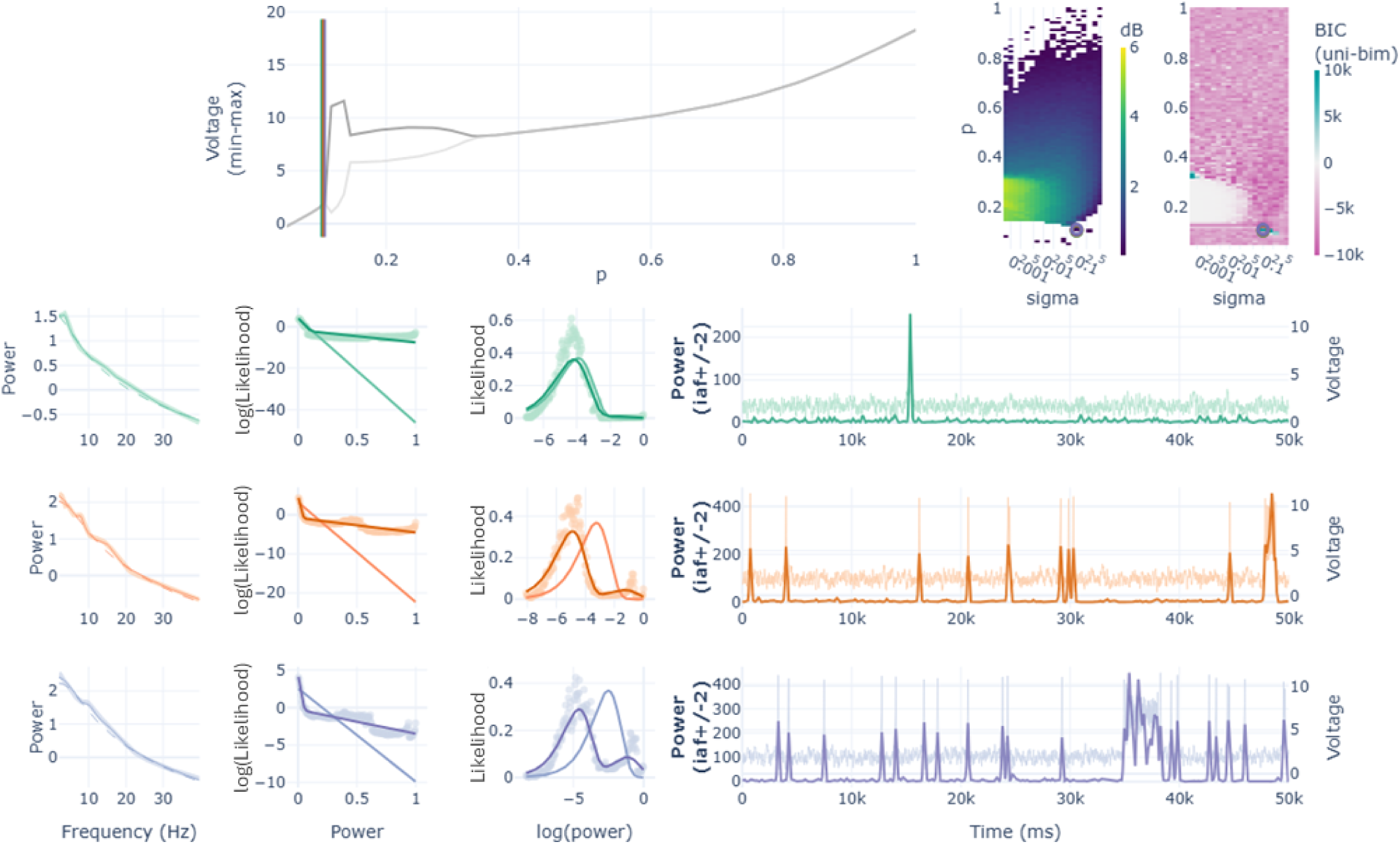
Samples of bimodalities found in single node simulations (Fig. 4) at the saddle node bifurcation (p=[0.1025, 0.105, 0.1075] with high noise (sigma=0.1). These bimodalities represent the switching behaviour of the node between the fixed point state and the limit cycle. Note that increasing p, rises the frequency of switching. All samples fitted better bimodal exponentials: green trace [BIC: unimodal = −291663.23; bimodal = −304581.65], orange trace [BIC: unimodal = −224595.13; bimodal = −316788.16], and blue trace [BIC: unimodal = −151996.88; bimodal = −234628.44].

**Fig 6.1.**
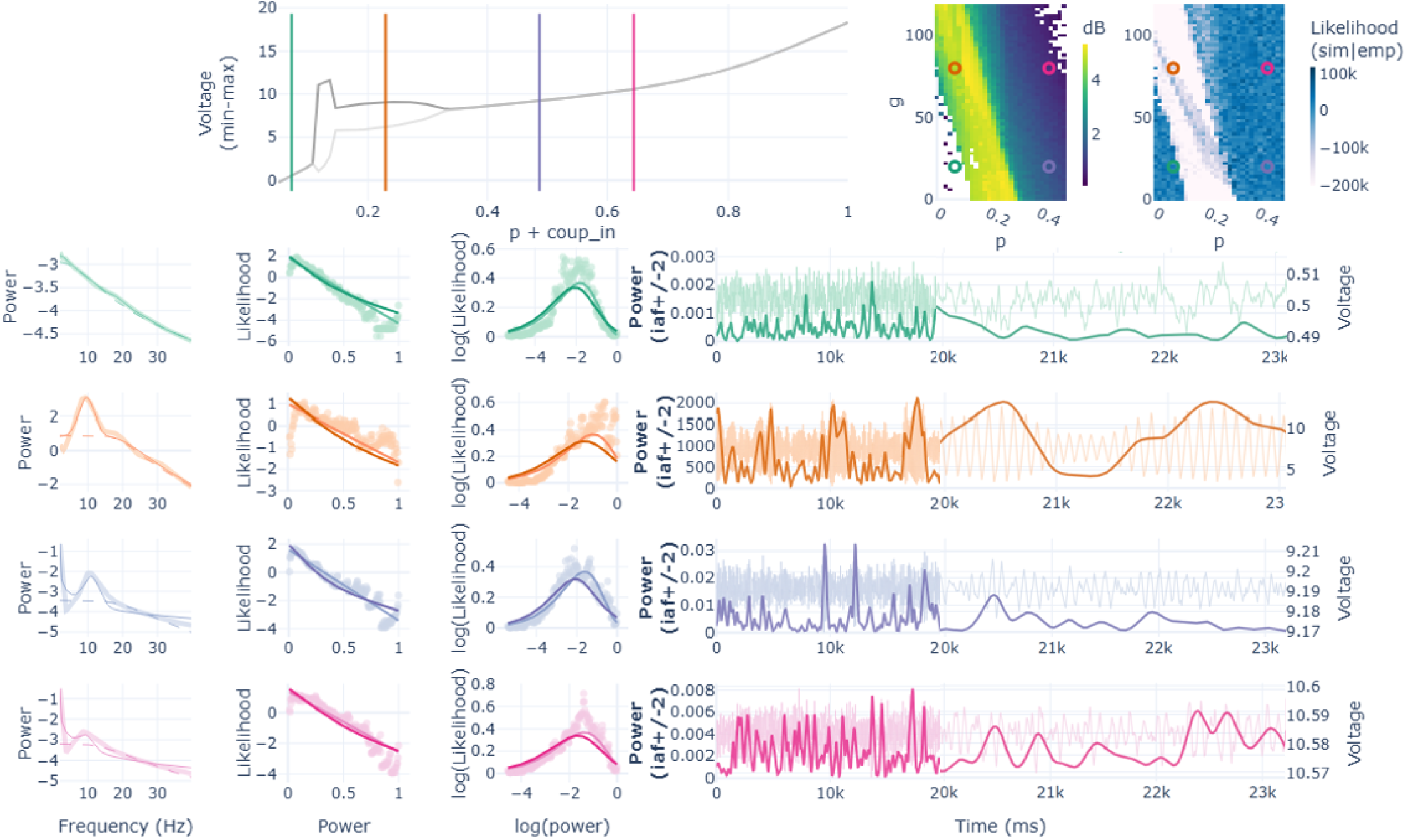
Samples of the network simulations for one region (V1 L). First row shows the bifurcation and two parameter spaces as reference for the picked simulations (in coloured vertical lines and open circles). Next rows show: 1) the power spectrum calculated from the data (thick line), the modeled spectra (thin line), and the modeled aperiodic component (dashed line); 2) the exponential function in two different coordinate spaces (i.e, Linear-Log and Log-Linear) with the scatter representing the histogram of the simulated TFR(*α*) and the lines representing both the unimodal (light colour) and the bimodal exponential models (dark colour); 3) the raw signal (light colour) and the TFR(*α*) values (dark colour) in time. The used parameters were (p, g) = [green (0.07, 20); orange (0.07, 80), blue (0.44, 20), pink (0.44, 80)].

**Supp. Table 1.**
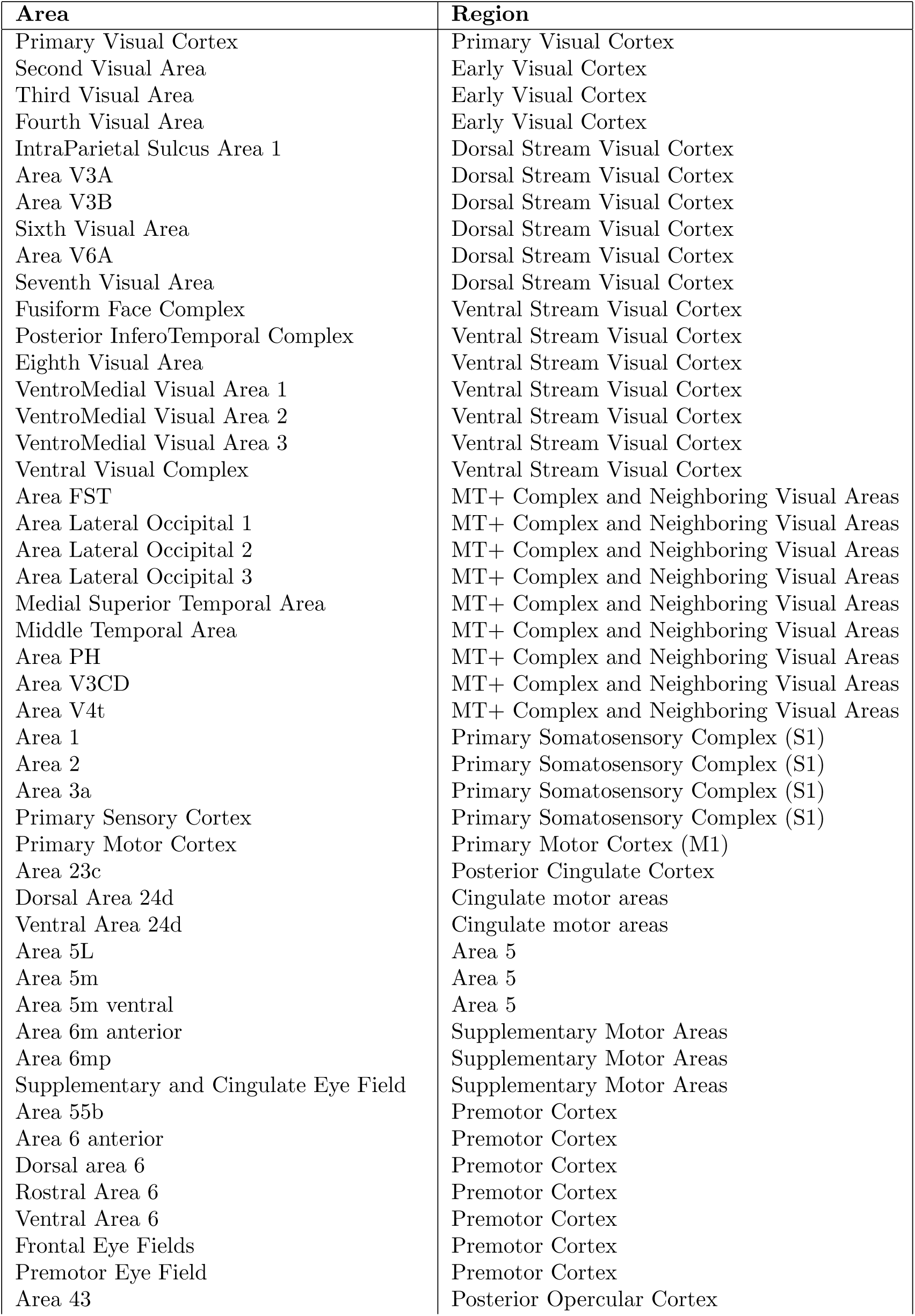
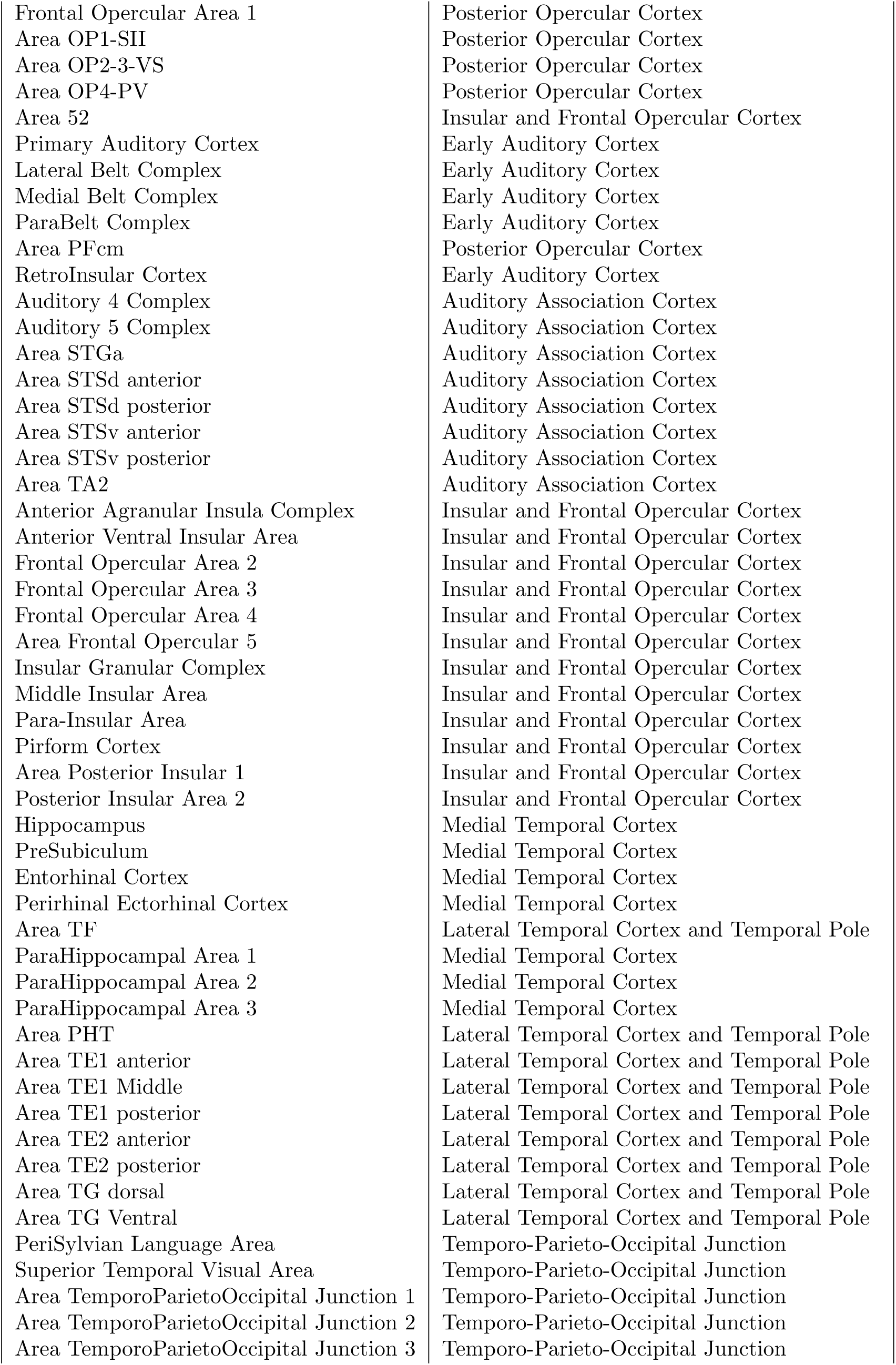
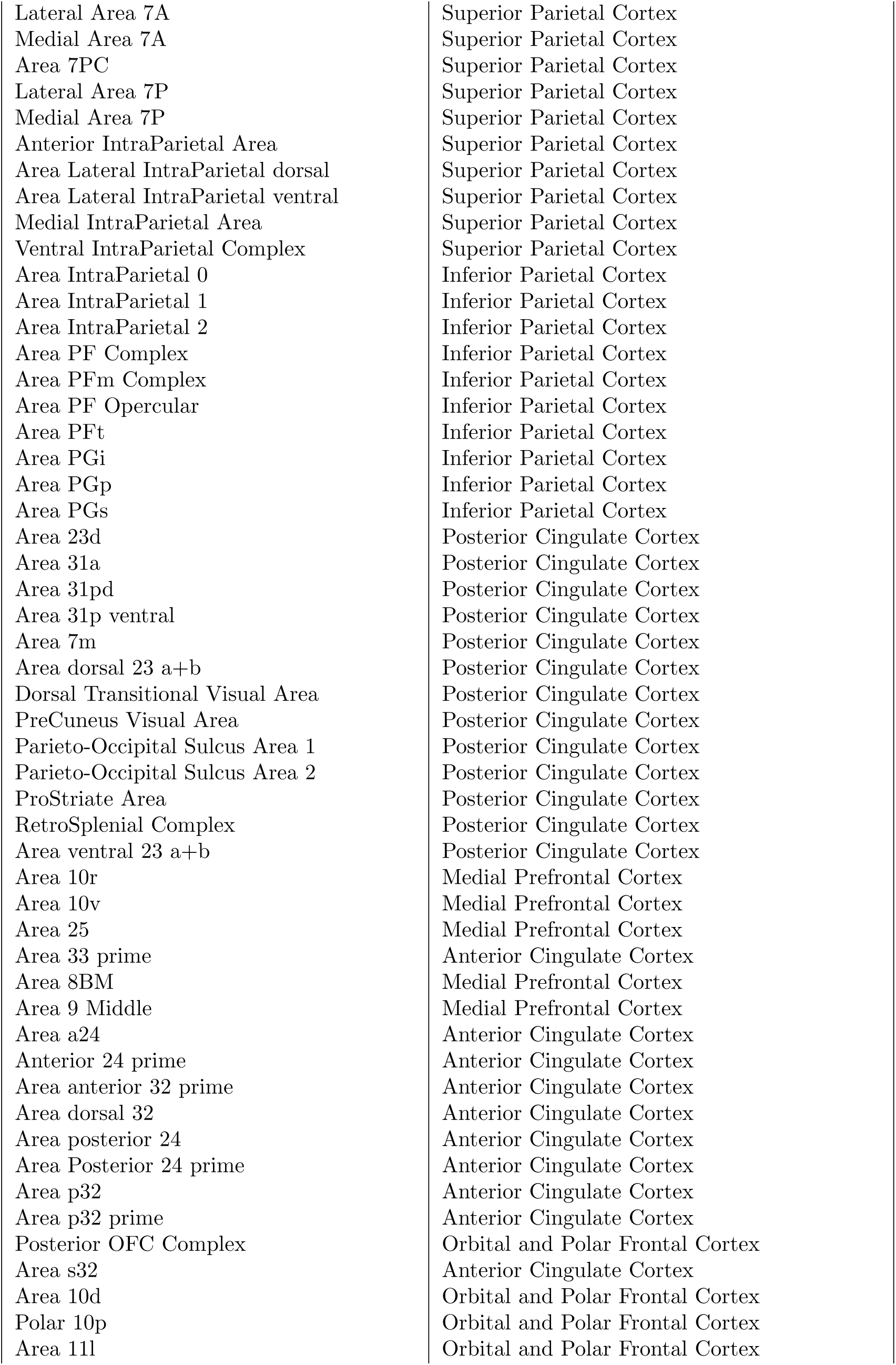
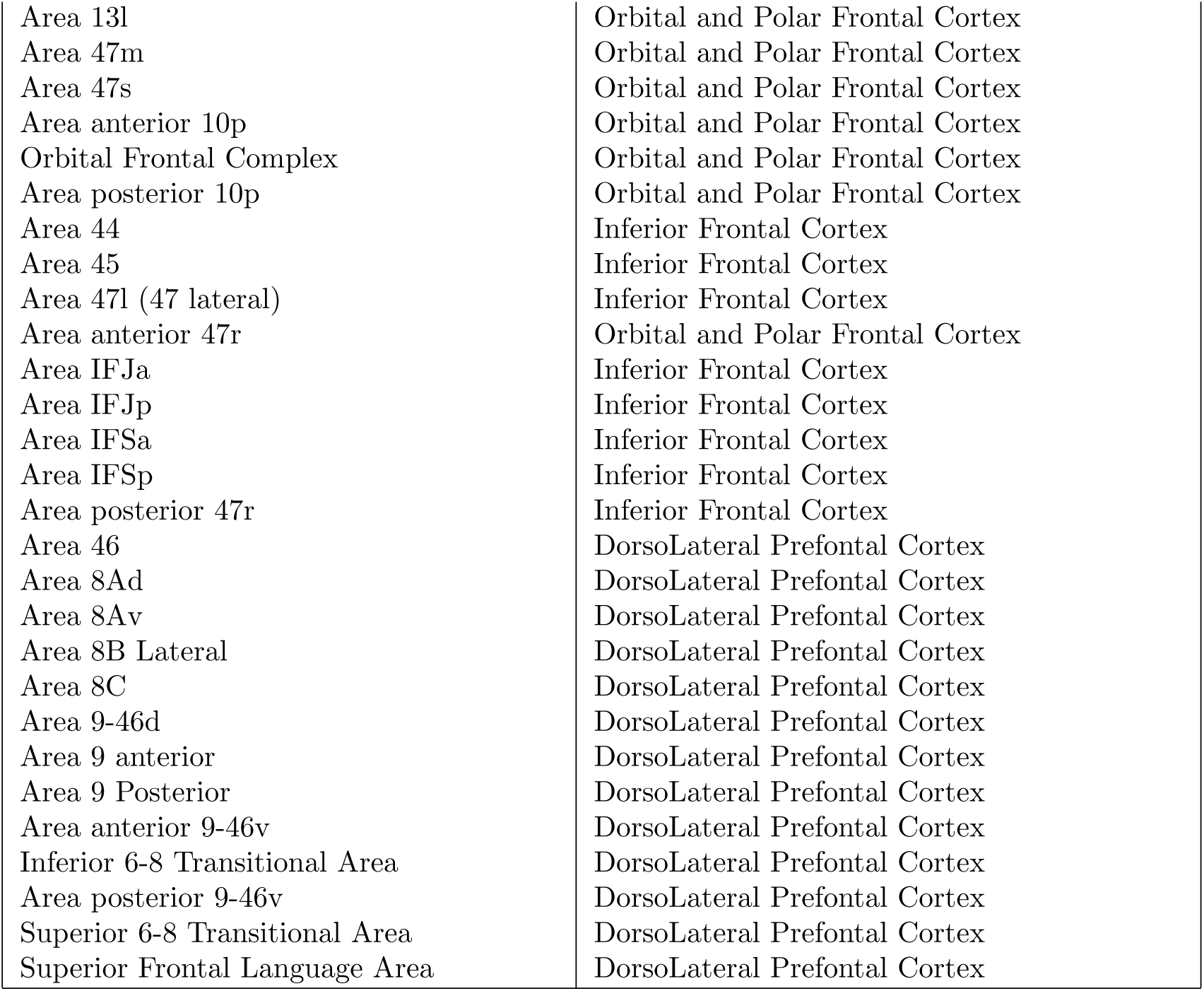
Areas of the HCPex atlas and corresponding Region groups (Huang et al., 2021).

**Supp. Table 2.**
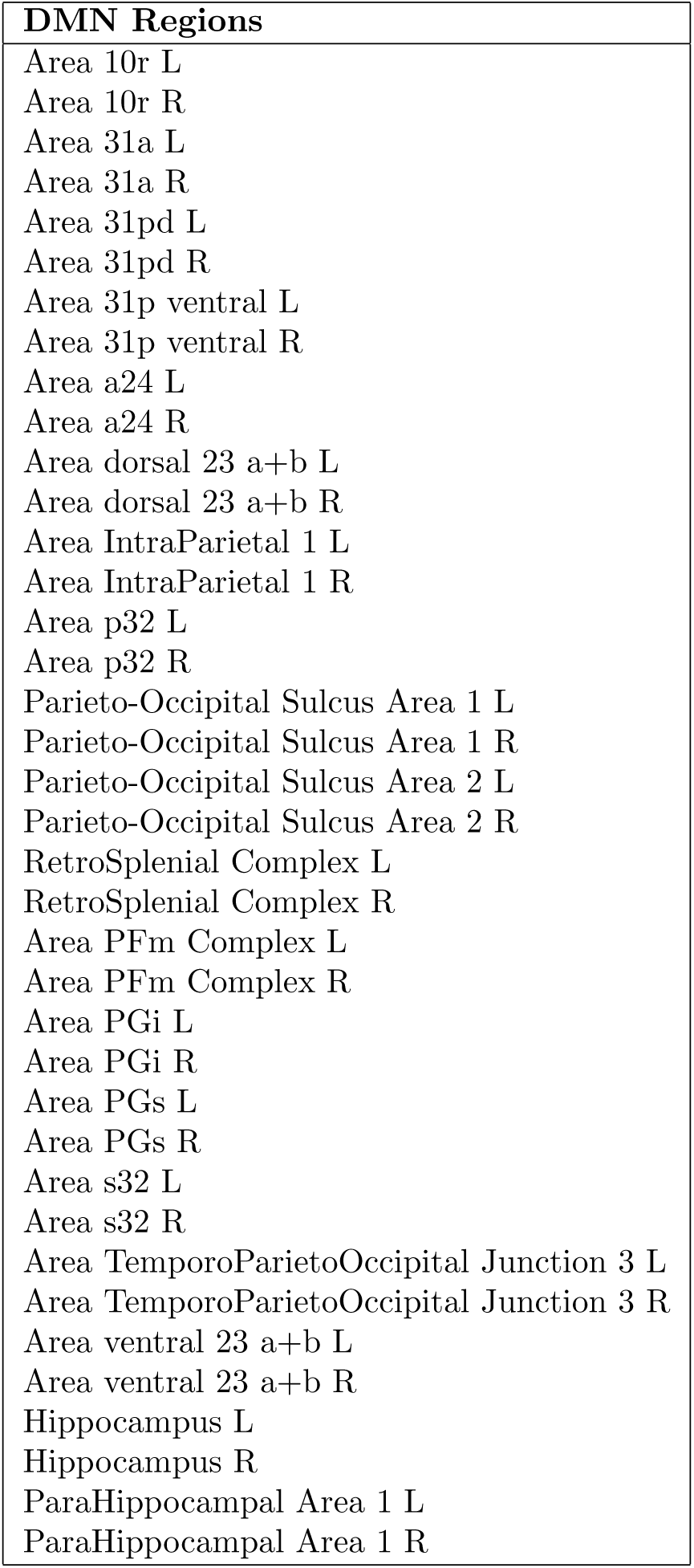
Brain regions of the HCPex atlas (Huang et al., 2021) included in the Default Mode Network (DMN) following Sandhu et al. (2020).

